# Influence of HIV-1 genomic RNA on the formation of Gag biomolecular condensates

**DOI:** 10.1101/2023.02.23.529585

**Authors:** Anne Monette, Meijuan Niu, Rebecca Kaddis Maldonado, Jordan Chang, Gregory S. Lambert, John M. Flanagan, Alan Cochrane, Leslie J. Parent, Andrew J. Mouland

## Abstract

Biomolecular condensates (BMCs) play an important role in the replication of a growing number of viruses, but many important mechanistic details remain to be elucidated. Previously, we demonstrated that pan-retroviral nucleocapsid (NC) and the HIV-1 pr55^Gag^ (Gag) proteins phase separate into condensates, and that HIV-1 protease (PR)-mediated maturation of Gag and Gag-Pol precursor proteins yield self-assembling BMCs having HIV-1 core architecture. Using biochemical and imaging techniques, we aimed to further characterize the phase separation of HIV-1 Gag by determining which of its intrinsically disordered regions (IDRs) influence the formation of BMCs and how the HIV-1 viral genomic RNA (gRNA) could influence BMC abundance and size. We found that mutations in the Gag matrix (MA) domain or the NC zinc finger motifs altered condensate number and size in a salt-dependent manner. Gag BMCs were also bimodally influenced by the gRNA, with a condensate-promoting regime at lower protein concentrations and a gel dissolution at higher protein concentrations. Interestingly, incubation of Gag with CD4^+^ T cell nuclear lysates led to the formation of larger BMCs as compared to much smaller ones observed in the presence of cytoplasmic lysates. These findings suggests that the composition and properties of Gag-containing BMCs may be altered by differential association of host factors in nuclear and cytosolic compartments during virus assembly. This study significantly advances our understanding of HIV-1 Gag BMC formation and provides a foundation for future therapeutic targeting of virion assembly.

## Introduction

Recent investigations have focused attention on the study of membraneless organelles known as biomolecular condensates (BMCs) due to their implicated role in the control and compartmentalization of a growing number of cellular processes in both nuclear and cytoplasmic compartments [1]. BMC formation is promoted by multivalent interactions between proteins and nucleic acids, driven in part by the presence of intrinsically disordered regions (IDRs) in proteins leading to liquid-liquid demixing [2]. Liquid-liquid phase separation (LLPS) relies on the coordinated condensation of proteins and nucleic acids into BMCs, driving compartmentalization, spatial organization, and coordinated biological functions [3]. BMCs are involved in many cellular processes that require concentration of protein-nucleic acid complexes for function [4], including DNA repair [5], transcription initiation/elongation [6, 7], and mRNA translation [8, 9]. Aberrant or less liquid-like BMCs generated by pathological gene mutations or exposure to metals are proposed to cause neurodegenerative diseases [10-13], and more recently, tumorigenesis [14, 15]. Recent studies have also indicated that BMCs may influence anti-cancer drug efficacy due to differential drug concentrations within condensates [16, 17].

BMCs appear to be fundamental to the replication cycle of several viruses [18]. Widely varying terms have historically been used to refer to viral BMCs, including viral replication compartments, viral ribonucleoprotein complexes, virosomes, virus factories, viroplasm, mini-organelles, inclusion bodies, Negri bodies, and liquid organelles [19-28]. *In silico* prediction and meta-analyses have demonstrated disproportionately high frequency of IDRs in viral proteins relative to those of eukaryotes, with a notably high preponderance in retroviral Gag proteins [29, 30]. Viral protein IDRs are speculated to facilitate multivalent interactions, conferring a competitive advantage over host protein for their intracellular replication [31].

The HIV-1 Gag protein orchestrates virion assembly by binding to viral genomic RNA (gRNA), undergoing Gag-Gag multimerization, and recruiting host factors involved in the budding process [32-34]. Our previous work demonstrated that Gag intracellular complexes exhibit fusion and fission events characteristic of BMCs [35]. HIV-1 Gag also contains IDRs juxtaposed with RNA-binding domains (RBDs) [13]. Both IDRs and RBDs promote sequestration of protein and RNA partners in dynamic BMCs and confer liquid-like behavior during virus assembly [36, 37]. During HIV-1 virion maturation, Gag is proteolytically cleaved into the MA, capsid (CA), NC, and p6 proteins, and Gag-Pol is processed into the viral enzymes reverse transcriptase (RT), integrase (IN), and PR [32]. Analysis of the HIV-1 structural proteins revealed a high degree of disorder, which may not only contribute to formation of BMCs but also provide antigenic variability and immune evasion, complicating vaccine design [38, 39].

The highly basic and disordered NC domain in Gag binds gRNA via its two CCHC zinc finger (ZnF) domains conferring RNA-binding specificity, nucleic acid chaperone activity, and RNA packaging activity [40-43]. Our earlier work demonstrated that the disordered HIV-1 NC protein undergoes zinc-dependent phase separation, regulating gRNA positioning and trafficking [44]. More recently, we demonstrated that HIV-1 NC, CA, RT, and IN proteins phase separate in proportion to their degrees of intrinsic disorder, and furthermore, that virus maturation by HIV-1 PR generates NC-initiated condensates driving maintenance of functional viral core BMCs [45]. We are also intrigued by the observation that the HIV-1 MA protein also possesses high levels of predicted intrinsic disorder [46]. Considering that both IDR-containing NC and MA domains of Gag interact with gRNA as a requirement for packaging, membrane targeting, and virus assembly, in the current work we investigated how their IDRs contribute to HIV-1 Gag BMC formation [47-51].

The Gag MA domain contains a bipartite membrane-binding domain consisting of an N-terminal myristic acid moiety and a highly basic amino acid region [52] that preferentially interacts with phosphatidylinositol-(4, 5)-bisphosphate (PIP_2_)-containing lipids at the plasma membrane [53-55]. To assess the contributions of basic residues in MA and the ZnFs of NC to Gag BMC formation, we performed *in vitro* biochemical and imaging experiments to address how MA and NC mutations affect the physical properties of HIV-1 Gag-gRNA BMCs. We observed that mutations in MA and NC differentially affected the propensity of Gag to phase separate with gRNA into liquid-like spherical droplets or gels. Furthermore, by incubating Gag with fractionated cell lysates, we observed that whereas cytoplasmic lysates promoted smaller, more numerous Gag condensates, nuclear lysates promoted much larger Gag condensates, indicating that host factors found in different cellular compartments may bind to Gag BMCs and influence their physical properties.

## Results

### The MA and NC RNA-binding motifs of HIV-1 Gag are disordered

A core requirement for replication and budding of infectious retrovirus particles is the specific binding of structural viral proteins to their cognate gRNA. In our previous work, we observed that retroviral NC proteins have prominent and conserved IDRs promoting phase separation via ZnF RNA-binding motif-dependent zinc (Zn^2+^) coordination [44]. We additionally discovered that like Gag, structural NC proteins from many other viruses also contain conserved IDRs juxtaposed by metal-regulated RNA-binding motifs [13]. Intrinsic disorder in RNA-binding proteins is advantageous to their function by providing flexibility permitting nucleic acid scanning, and the binding to a variety of different RNA sequences while also ensuring RNA-binding specificity [56, 57].

To determine whether RNA-binding IDRs in the HIV-1 MA and NC domains could influence the ability of Gag to phase separate with gRNA, we used PONDR and DisEMBL algorithms to assess overall disorder scores to map IDRs within the HIV-1 Gag protein [58, 59]. The full-length Gag protein was found to be 52.40% disordered overall, whereas the MA and NC subdomains were 59.09% and 43.64% disordered, respectively (Figure 1A). Interestingly, disorder was higher in both the RNA-binding MA highly basic region (MA-HBR) (i.e., 70.00%) [51] and RNA-binding NC ZnFs (48.57%). Although the NC domain of Gag contributes to phase separation and can itself produce large cytoplasmic aggregates within stress granules [44, 45, 60, 61], the MA domain has yet to be shown to contribute to phase separation of Gag. However, a number of studies have shown that MA forms trimers [62-64], and that MA alone can produce VLPs [65, 66], suggesting that, similar to NC, the mature MA protein participates in protein-protein interactions.

**Figure 1:**
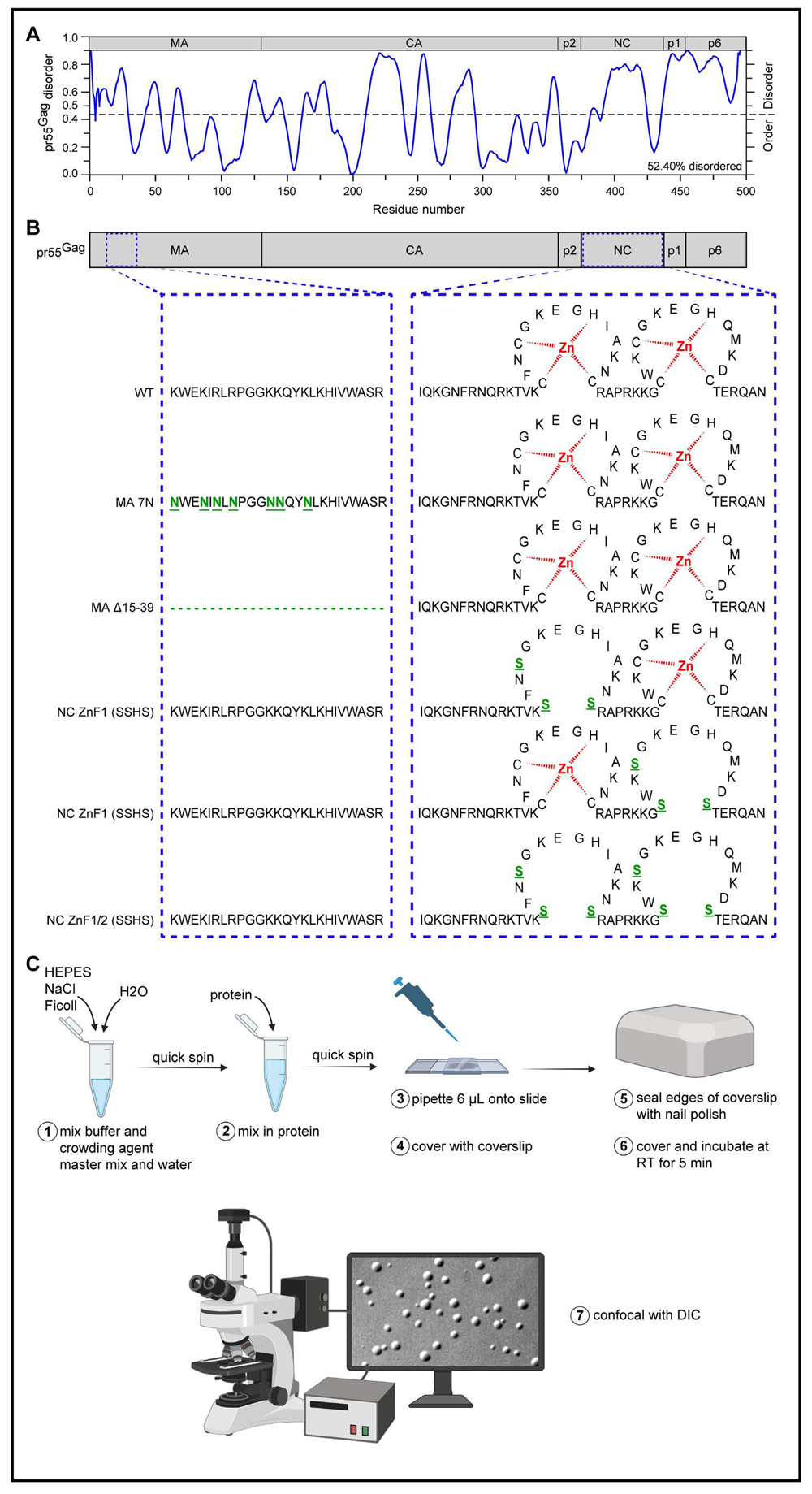
Investigations into disorder of the matrix and nucleocapsid viral RNA-binding motifs of HIV-1 Gag. **(A)** DisEMBL algorithm-generated depiction of disorder across HIV-1 Gag polyprotein (GenBank: AAC82593.1), demonstrating that the matrix (MA) RNA-binding region (MA-RBD) and nucleocapsid (NC) domains are inherently disordered. **(B)** Depiction of WT Gag, MA, and NC mutants used in this study. Mutations and/or deletions made to Gag are shown in green, and zinc coordination is shown in red. **(C)** Depiction of *in vitro* experimental method used in this study.

To determine the extent that MA and NC IDRs contribute to phase separation of Gag, we cloned, expressed, and purified HIV-1 Gag wild-type (WT) and mutant proteins. For MA mutants, positively charged lysine (Lys, K) and arginine (Arg, R) residues were substituted for uncharged asparagine (Asn, N; mutant MA 7N), or the entire MA highly basic region was deleted (i.e., MA Δ15-39) [52] (Figure 1B). For NC mutants, the Zn^2+^ coordinating cysteines (Cys, C) of CCHC ZnFs, which generate RNA-binding tertiary structure, were replaced with serine (Ser, S) to create three mutants: NC ZnF1 mutant (i.e., NC ZnF1 SSHS); NC ZnF2 mutant (i.e., NC ZnF2 SSHS); or NC ZnF1/ZnF2 double mutant (i.e., NC ZnF1/2 SSHS) [67, 68] (Figure 1B). To assess the effects of these amino alterations on the ability of Gag to phase separate, highly purified recombinant Gag proteins were mixed with buffers containing Ficoll-400 as a crowding agent [44, 69], transferred to slides, sandwiched with coverslips, and sealed for the visualization of condensate formation using confocal and differential interference contrast (DIC) microscopy (Figure 1C). Of note, proteins that interact with nucleic acids, in addition to those that oligomerize or aggregate, are prone to having low solubility potential during protein crystallization [70]. Therefore, ZnF-containing proteins such as HIV-1 Gag are typically purified using high salt buffers to promote their solubility [71]. In this work, all proteins were stored in 500 mM NaCl, with exception of the NC ZnF2 mutant and the NIH WT Gag proteins which were both stored at 1 M NaCl. These salt concentrations were considered to be the baseline that would maintain protein solubility, although the higher salt concentration could have represented a potential inherent limitation of the study. However, at the highest protein concentration tested (40 µM), the overall salt concentration was adjusted for as a result of the differences in protein concentrations normalized in the assay (e.g., NC ZnF1: NC ZnF2: NC ZnF1/2 = 384:442:389 mM NaCl). Thus, despite these small differences in overall salt concentrations, we observed similar patterns of phase separation for these proteins during the construction of phase diagram, with most prominent condensate formation for all three constructs at 40 µM protein in crowding buffer containing 150 mM NaCl (see below Figures 2-3), which was the same salt concentration used for the other experiments involving gRNA (see below Figure 6).

**Figure 2:**
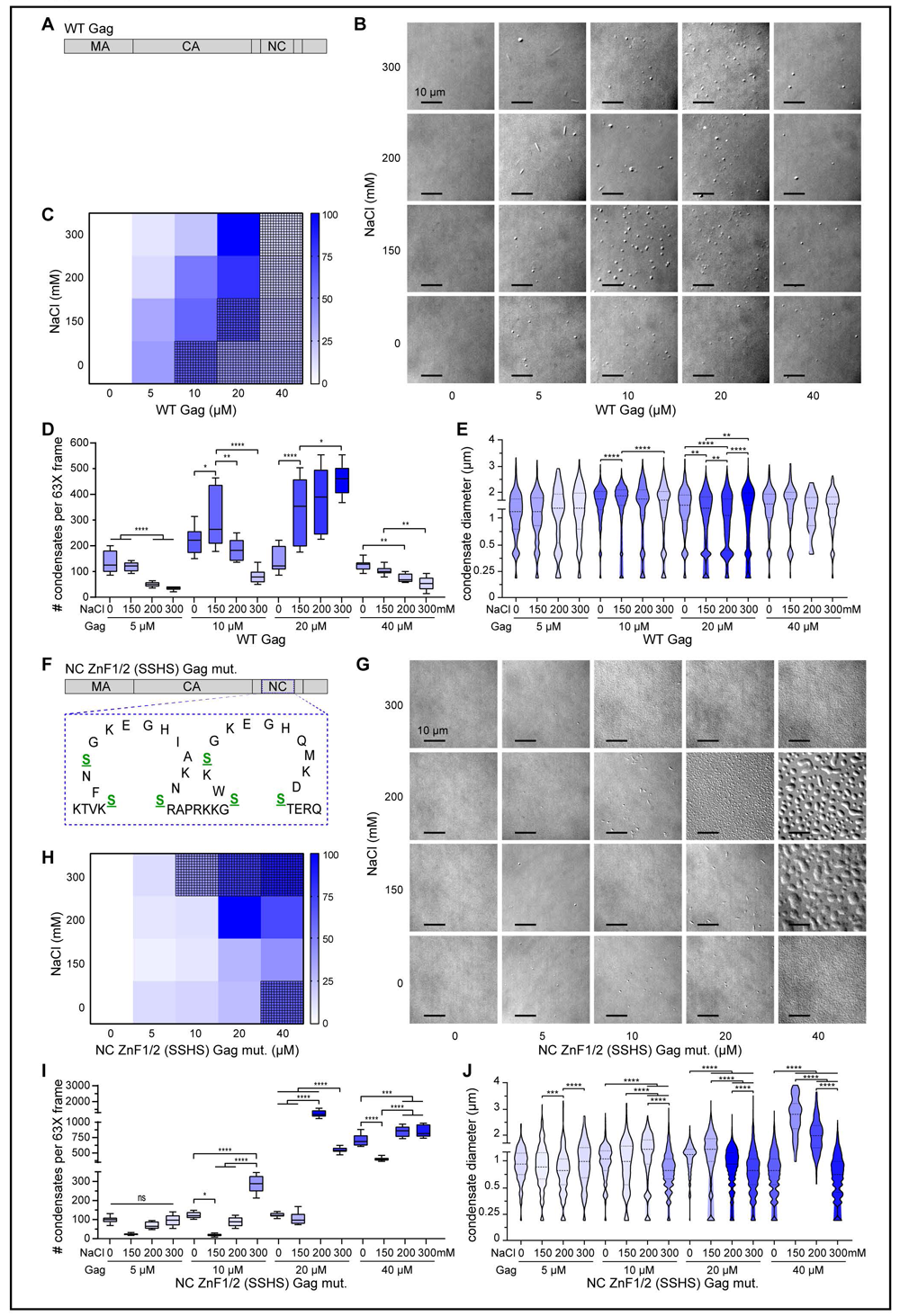
Effects of HIV-1 Gag double NC zinc finger (ZnF) mutations on phase diagrams and condensate sizes. Increasing concentrations of **(A-E)** full-length recombinant WT HIV-1 NL4-3 pr55 Gag or **(F-J)** the double Gag NC ZnF1 and ZnF2 mutant proteins (i.e., 0-40 μM) were mixed with crowding buffers composed of 20 mM HEPES, 150 mg/ml Ficoll-400, and increasing salt (NaCl, pH 7.4) concentrations (i.e., 0- 300 mM). (B and G) Images of phase separated condensates were captured using confocal microscopy and differential interference contrast (DIC), and statistical analyses were performed to create comparative phase diagrams in the form of heat maps (C and I), relative differences in condensate numbers (D and I) and condensate sizes (E and J). Heat maps represent number (#) of condensates per low-magnification frames. Grid patterns were then overlaid onto heat map squares to show experimental conditions in which proteins were observed to form gels. Box and whisker and violin plots were pseudocolored according to heat maps. Statistics represent *, p<0.05; **, p<0.01; ***, p<0.001; ****, p<0.0001, as derived from one-way ANOVA (with Tukey’s multiple-comparisons test) and 95% CI for multiple comparisons. mM, millimolar; µM, micromolar; µm, micron.

**Figure 3:**
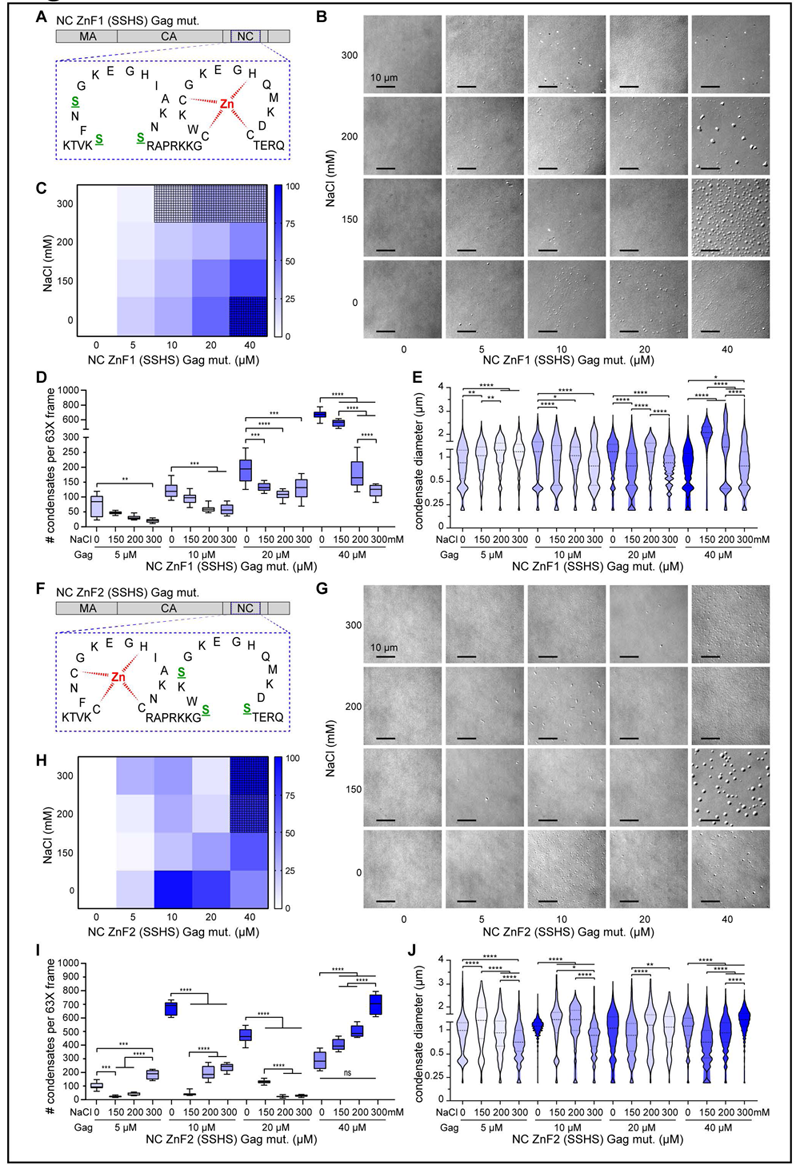
Effects of HIV-1 Gag single NC zinc finger mutations on phase diagrams and condensate sizes. Increasing concentrations of **(A-E)** full-length recombinant Gag NC ZnF1 mutant or **(F-J)** Gag NC ZnF2 mutant proteins (i.e., 0-40 μM) were mixed with crowding buffers composed of 20 mM HEPES, 150 mg/ml Ficoll-400, and increasing salt (NaCl, pH 7.4) concentrations (i.e., 0-300 mM). (B and G) Images of phase separated condensates were captured using confocal microscopy and differential interference contrast (DIC), and statistical analyses were performed to create comparative phase diagrams in the form of heat maps (C and I), relative differences in condensate numbers (D and I) and condensate sizes (E and J). Heat maps represent number (#) of condensates per 63 X magnification frames. Grid patterns were then overlaid onto heat map squares to show experimental conditions in which proteins were observed to form gels. Box and whisker and violin plots were pseudocolored according to heat maps. Statistics represent *, p<0.05; **, p<0.01; ***, p<0.001; ****, p<0.0001, as derived from one-way ANOVA (with Tukey’s multiple-comparisons test) and 95% CI for multiple comparisons. mM, millimolar; µM, micromolar; µm, micron.

### HIV-1 MA and NC mutations induced differential changes in Gag condensation

Our past work demonstrated that, as observed for recombinant HIV-1 NC protein, *in vitro* HIV-1 Gag condensates could be dissolved by Zn^2+^ chelation or ejection [44]. Therefore, we expected that the Zn-chelating Cys-His boxes in the NC domain of Gag were primarily responsible for condensate formation. This notion was supported by our most recent observation that in cells, the HIV-1 NC protein serves as a scaffold onto which other viral proteins condense to form HIV-1 BMCs [45]. We therefore performed experiments to generate phase diagrams for each recombinant Gag protein by assessing the effect of crowding buffer containing additional NaCl with increasing protein concentrations on condensate formation (ranging from 0 to 40 µM protein; 0 to 300 mM NaCl). As previously reported, WT Gag optimally phase separated into spherical condensates between concentrations of 10 - 20 µM at 150 mM NaCl (Figure 2B, D), with sizes ranging from 0.5 to 2 μm in diameter [44, 72] (Figure 2B, E). At higher viral protein concentrations [45] and increasing salt concentrations, Gag formed a gel (Figure 2B, C). At lower protein concentrations and higher salt concentrations, Gag formed rod-like structures (Figure 2B). In previous studies, recombinant HIV-1 Gag proteins self-assemble into spherical structures *in vitro* [73-75], while HIV-1 and RSV CA-NC proteins assemble into rod-like structures [76]. Although no consensus has yet been reached on the mechanism by which purified recombinant Gag or newly-translated Gag proteins assemble in cell-free systems [77], disordered regions of Gag have been demonstrated to play essential roles in its multimerization [78].

Next, we tested the ability of the Gag NC ZnF double mutant to form condensates *in vitro* (Figure 2F). Distinct NC ZnF1/2 SSHS mutant condensates were most numerous at increasing protein (20-40 µM) and salt (150-200 mM) concentrations (Figure 2G, I). Gel formation by the double NC ZnF mutant was visible at higher protein and salt (300 mM) concentrations (Figure 2G, H). This mutant protein exhibited a leftward shift in phase diagrams compared to the WT (Figure 2C, H), suggesting that higher salt concentration was required to accelerate condensate and gel formation for the NC ZnF double mutant. Furthermore, the double mutant displayed gel formation at both low and high salt concentrations, suggesting that it is more susceptible to re-entrant phase separation dynamics (i.e., monotonic variation of a single thermodynamic control parameter driving proteins from a phase-separated state to a macroscopically similar state via two-phase transitions) [79, 80], possibly resulting from the loss of the NC domain’s condensate scaffolding activity [45]. Interestingly, at the highest protein (40 µM) and middle salt (150-200 mM) concentrations, double NC mutant condensates were much larger and less spherical (Figure 2G, J), merging into larger droplets that were flatter then condensates at the 150 mM salt concentration (Figure 2G). Condensate size scaling is governed by Ostwald ripening (i.e., smaller particles in solution dissolve onto larger particles to reach a more thermodynamically stable state and minimizing surface to area ratio), coalescence and interfacial tension [81]. Based on these principles, our observations suggest that the NC double mutant attained a less crystal-like structure and favoured a more deformed liquid-or fluid-like state. The increased liquid-like appearance could result from diminished surface tension as a consequence of short-range positional order [82] possibly due to modified scaffold to client:surfactant ratios [83] or diminished interaction strength between IDRs [84].

Next, we examined the single NC ZnF1 and ZnF2 mutants (Figure 3A, F). Like the double NC mutant, the highest number of condensates were formed at high protein (40 µM) and salt concentrations (Figure 3B-D and 3G-I). Increasing salt concentration for these single mutants also resulted in the formation of larger condensates (Figure 3B,E and 3G-J) with apparent diminished sphericity observed for the NC ZnF1 mutant (Figure 3B) at 40 µM protein and 150-200 mM salt concentrations, similar to the NC double mutant (Figure 2G). Although both NC single mutants exhibited an upward shift in their phase diagrams (Figure 3C, H), the NC ZnF1 mutant displayed a re-entrant gelling feature (Figure 3B-C), similar to the NC double mutant (Figure 2G-H). The NC ZnF2 mutant condensates were more spherical at the 40 µM protein and 150 mM salt concentrations (Figure 2M), whereas the NC ZnF1 mutant condensates were more abundant and less spherical (Figure 3B). In addition, NC ZF1 droplets at 40 µM protein and 150 mM NaCl appeared to have undergone fusion or were coalescing, indicative of a more fluid-like state, as observed for the double NC mutant (Figure 2G). Although both NC single mutants presented features common to the NC double mutant, including increased gel formation at higher salt concentrations, the NC ZnF1 mutant phase diagram had closer resemblance to the properties of the double mutant (Figure 3B and 2G). Based on these observations, NC ZnF1 appeared to make a larger contribution to condensate formation than NC ZnF2. These differences in ZnF1 and ZnF2 may be attributed to their independent functions within Gag, in which ZnF1 is needed for correct core morphology, proviral cDNA synthesis [85, 86], specific binding to psi [87], and gRNA selection and packaging [88], whereas ZnF2 nucleates nucleic acid association, contributes to Gag-gRNA complex stabilization [89], and influences localization to the plasma membrane (see Table 1) [90].

We next examined the effects of Gag MA mutants on condensate formation (Figure 4A, F). Interestingly, both 7N and Δ15-39 MA mutants formed similar rod-like structures at high protein and low salt concentrations analogous to those formed by CA-NC (Figure 4B, G). The MA mutants also formed higher overall numbers of condensates (Figure 4B, D, G, I) and the Δ15-39 MA mutant exhibited gel formation at higher protein and salt concentrations (Figure 4G-H). However, despite minimal modifications to the phase diagrams upon mutations of MA (Figure 4C, H), the relative quantity, morphology, sphericity, surface tension, size and homogeneity of the condensates [72], like WT Gag, remained optimal at 10-20 µM protein and 150-200 mM salt concentrations (Figure 4B, D, E, G, I, J). Taken together, these results suggested that mutations made to the MA domain had a less prominent effect on Gag condensation than those made to the NC domain.

**Figure 4:**
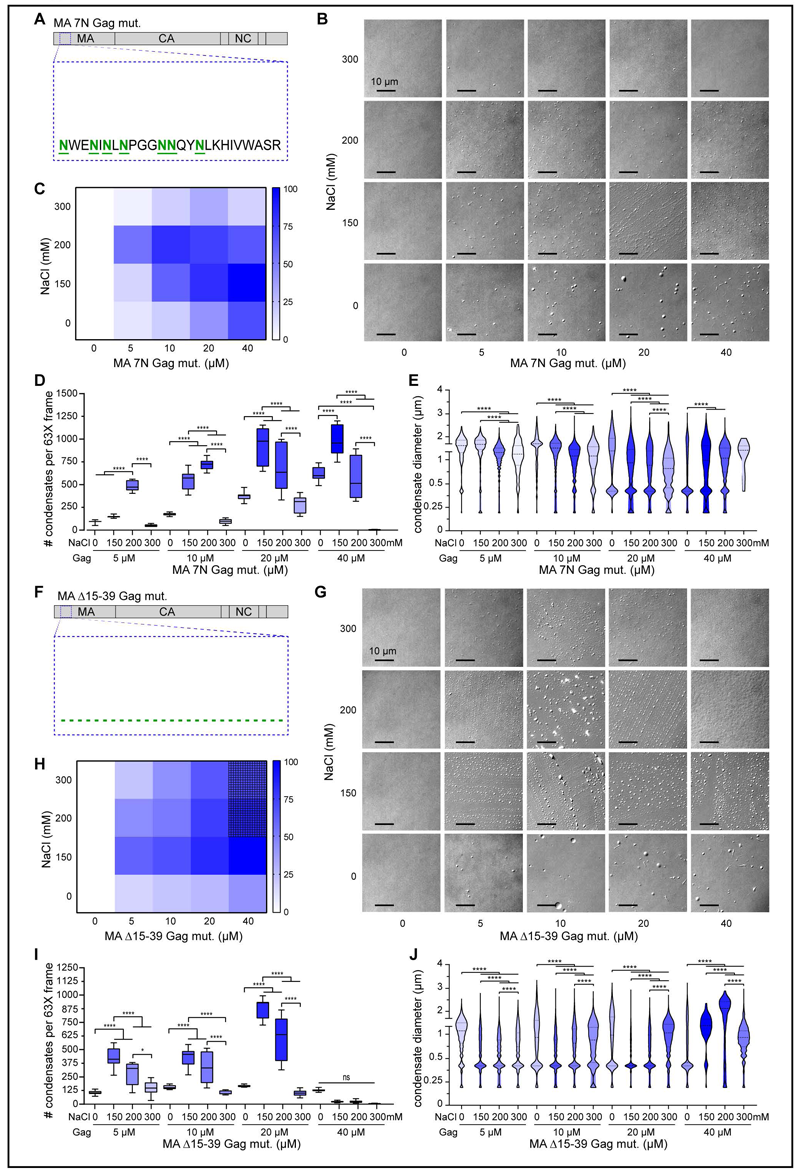
Effects of HIV-1 Gag MA substitution mutations or RBD deletion on phase diagrams and condensate sizes. Increasing concentrations of **(A-E)** full-length recombinant MA 7N Gag mutant or **(F-J)** MA Δ15-39 Gag mutant proteins (i.e., 0-40 μM) were mixed with crowding buffers composed of 20 mM HEPES, 150 mg/ml Ficoll-400, and increasing salt (NaCl, pH 7.4) concentrations (i.e., 0-300 mM). (B and G) Images of phase separated condensates were captured using confocal microscopy and differential interference contrast (DIC), and statistical analyses were performed to create comparative phase diagrams in the form of heat maps (C and I), relative differences in condensate numbers (D and I) and condensate sizes (E and J). Heat maps represent number (#) of condensates per 63 X magnification frames. Grid patterns were then overlaid onto heat map squares to show experimental conditions in which proteins were observed to form gels. Box and whisker and violin plots were pseudocolored according to heat maps. Statistics represent *, p<0.05; **, p<0.01; ***, p<0.001; ****, p<0.0001, as derived from one-way ANOVA (with Tukey’s multiple-comparisons test) and 95% CI for multiple comparisons. mM, millimolar; µM, micromolar; µm, micron.

### HIV-1 gRNA induced a bimodal shift in Gag condensation

We recently described a high degree of conservation of NC ZnF-embedded IDRs across retroviral Gag proteins that affected its condensation which, in turn, regulated positioning and trafficking of the gRNA [44]. We also reported that nuclear Gag colocalizes with unspliced vRNA in discrete ribonucleoprotein complexes [91], suggesting that the vRNA may influence Gag BMC formation. We therefore performed experiments using different ratios of Gag to gRNA purified from virions to measure the effect of gRNA on Gag condensation *in vitro*.

For these experiments, we used Gag:gRNA ratios similar to those found in virions (i.e., each immature virion contains ∼2400 Gag monomers and 2 copies of gRNA [92]. At 10 µM Gag concentration, 4 nM of purified gRNA represents approximately one copy per 2400 Gag molecules, whereas 8 nM gRNA represents approximately two copies per 2400 Gag molecules (Figure 5A). Interestingly, addition of gRNA to the *in vitro* droplet assay led to a biphasic shift in Gag phase diagrams. At a low Gag concentration (i.e., 2.5 µM), 8 nM gRNA (i.e., 8 gRNA:2400 Gag molecules) induced Gag condensation and increased particle size (Figure 5B-D), whereas at high Gag concentrations (i.e., 20-40 µM), increasing amounts of gRNA resulted in the dissipation of Gag gels (i.e., 16 nM gRNA:40 µM Gag reflects 1 gRNA:2400 Gag) (Figure 5B, C). This finding suggests that gRNA has a modulatory effect on Gag-gRNA BMCs, stimulating their formation at low protein concentration and causing re-entry from gel to liquid phase at high RNA concentrations. Similar to this finding, the influence of RNA on multiphasic condensate morphologies was recently reported [93-96]. In another report, RNA influenced condensate switching from one of Gag intra-protein IDR-RNA-binding motif interactions to one of RNA-binding motif-RNA interaction, with low RNA concentrations inducing condensates and high RNA concentrations dissolving or causing the demixing of condensates [97].

**Figure 5:**
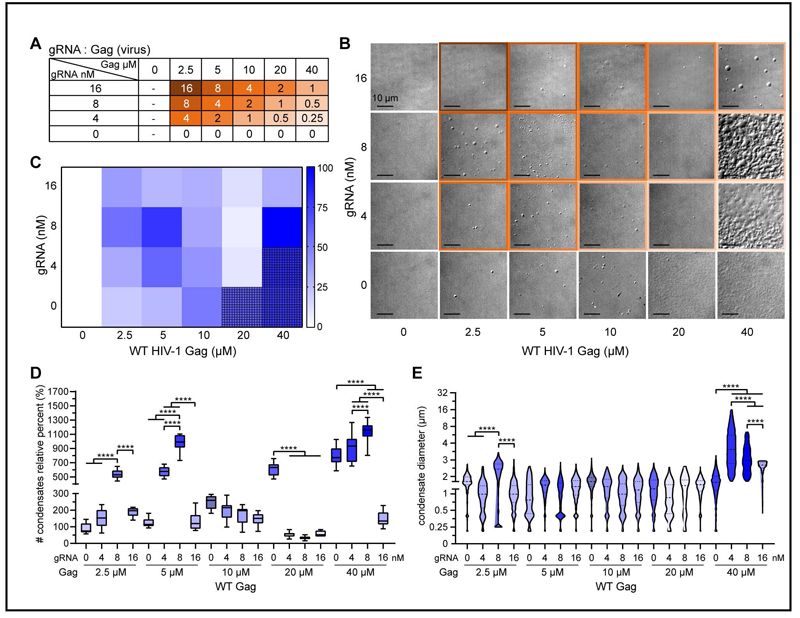
WT NL4-3 HIV-1 vRNA influences phase separation of HIV-1 pr55 Gag protein. **(A)** HIV-1 vRNA to 2400 Gag ratios tested, where 1 reflects one vRNA copy per virus **(B)** Increasing concentrations of full-length recombinant HIV-1 NL4-3 pr55 Gag protein (i.e., 0-40 μM) were mixed with crowding buffers composed of 20 mM HEPES, 150 mg/ml Ficoll-400, and 150 mM NaCl, pH 7.4, and in increasing concentrations of HIV-1 NL4-3 vRNA purified from viruses isolated from supernatants of HEK 293T cells transfected by pNL4-3. Frames surrounding DIC images are extrapolated from (A). **(C)** Heat map corresponding to phase diagram from panel (B), as calculated from number (#) of condensates per low-magnification frame, and demonstrating that HIV-1 vRNA induces a biphasic distribution of Gag condensates, where it induces Gag condensates at low protein concentration and dissipates Gag condensates at high protein concentration. Grid patterns placed over squares of heat map demarcate observed gelification of protein. **(D)** Box and whisker plot corresponding to data from (B) and pseudocolored according to (C). **(E)** Violin plots corresponding to sizes of condensates calculated from data presented in (B), and pseudocolored according to (C). Statistics represent ****, p<0.0001, as derived from one-way ANOVA (with Tukey’s multiple-comparisons test) and 95% CI for multiple comparisons. nM, nanomolar; µM, micromolar; µm, micron.

### Mutations in Gag MA and NC domains induced differential shifts in Gag-gRNA co-condensation

Guided by our observation that mutations within the gRNA-binding domains of HIV-1 Gag affected condensate formation (Figures 2-4), combined with the finding that 8 nM of gRNA produced the most striking effects on Gag phase separation at the 2.5 and 40 µM protein concentrations (Figure 5), we assessed the effect of mutations in MA and NC on condensation in the context of gRNA. The MA Δ15-39 and the NC ZnF1&2 mutants each exhibited an increased number of condensates at low protein concentration (Figure 6A, B). Interestingly, at 2.5 µM protein concentration, the NC ZnF mutants demonstrated increased condensate size, whereas a decrease in sphericity was observed for the MA mutant condensates upon addition of gRNA (Figure 6A-C). At 40 µM protein concentration, the presence of gRNA caused the MA mutants to form larger, non-spherical condensates resembling those of WT Gag (Figure 6D-F), suggesting that the gRNA promoted condensate formation. For the NC mutants tested at 40 µM, addition of gRNA caused an increase in condensate number and a decrease in condensate size (Figure 6D-F). Together, these data support a model in which the MA and NC domains of HIV-1 Gag are important modulators of condensate properties due to their ability to differentially interact with the gRNA.

**Figure 6:**
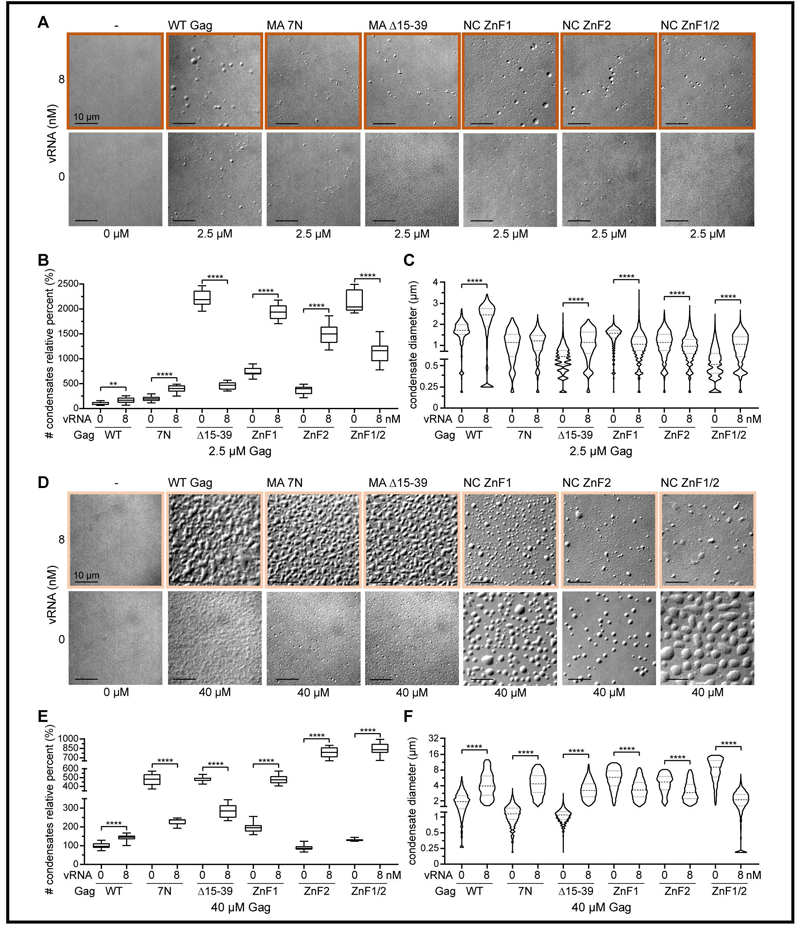
WT NL4-3 HIV-1 gRNA differentially influences phase separation of Gag MA and NC mutant proteins. **(A)** 2.5 μM of WT or mutant Gag proteins were mixed with crowding buffers composed of 20 mM HEPES, 150 mg/ml Ficoll-400, and 150 mM NaCl, pH 7.4, and either none or 8 nM of NL4-3 virus purified gRNA, for imaging using confocal imaging and DIC. Resulting images were analyzed for condensate numbers, as represented by a corresponding box and whisker plot **(B),** and for sizes, as represented by a corresponding violin plot **(C)**. **(D)** 40 μM of WT or mutant Gag proteins were mixed with crowding buffers composed of 20 mM HEPES, 150 mg/ml Ficoll-400, and 150 mM NaCl, pH 7.4, and either none or 8 nM of NL4-3 virus purified gRNA, for imaging using confocal imaging and DIC. Resulting images were analyzed for condensate numbers, as represented by a corresponding box and whisker plot **(E),** and for sizes, as represented by a corresponding violin plot **(F)**. Colored frames in panels (A) and (D) are representative of gRNA to Gag ratios presented in Figure 5. Statistics represent **, p<0.01; ****, p<0.0001, as derived from one-way ANOVA (with Tukey’s multiple-comparisons test) and 95% CI for multiple comparisons. mM, millimolar; µM, micromolar; µm, micron.

### Three-dimensional rendering of HIV-1 Gag-genomic RNA co-condenses

Based on our observation that high concentrations of gRNA disrupted the gelling of HIV-1 Gag (Figures 4 and 5), we examined where the gRNA was positioned within these BMCs. To visualize the BMCs in three-dimensions, we performed experiments using high concentrations of fluorescently-labeled Gag and gRNA, imaged a z-series of BMCs using confocal microscopy, and performed reconstructions [45]. In the absence of gRNA, we observed a gel of Gag-AF594 (red) at 30 µM (Figure 7A). Introducing 16 nM of gRNA (green) (i.e., 3 gRNA: 2400 Gag) transformed the Gag gel into spherical Gag condensates (Figure 7B). The gRNA also co-condensed and colocalized with the Gag condensates (Figure 7B), suggesting that the gRNA was responsible for dissolving Gag gels into spherical condensates. In the analyses of three-dimensional images, we observed that colocalization of the gRNA and Gag was limited to the core of these co-condensates, but the gRNA coated the exterior of the particles where it did not colocalize with Gag (Figure 7C). In examining the x-z planes (Figure 7D) and surface renderings of the Gag-gRNA complexes (Figure 7E), we observed that the gRNA appeared to sit on top of the Gag gel, potentially contributing to dissolution of the gel into individual BMCs.

**Figure 7:**
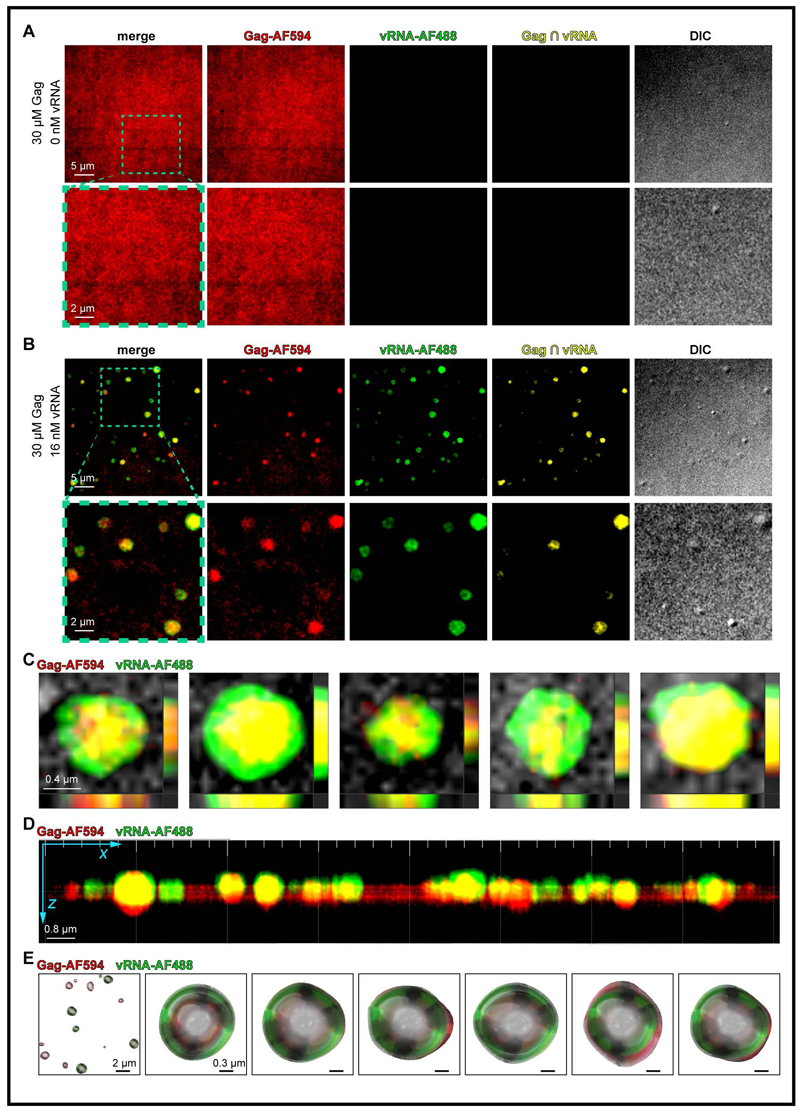
HIV-1 gRNA dissolves Gag gels into phase separated condensates. **(A)** Gag-AF5494 was mixed at 30 µM with crowding buffers composed of 20 mM HEPES, 150 mg/ml Ficoll-400, and 150 mM NaCl, pH 7.4, and was imaged using confocal microscopy and differential interference contrast **(B)** Same as in (A), but with the addition of 16 nM of vRNA purified from isolated WT NL4-3 virus that was green-labeled using fluorescence in situ hybridization, demonstrating that the vRNA dissolves red Gag gels into condensates and that these colocalize. **(C)** Single particle analyses provide evidence that the vRNA can be found within, and also outside of Gag condensates, as is also shown using Z-stack slice analyses **(D)**, providing evidence of interruption of Gag gel by surface vRNA association with Gag condensates. **(E)** 3 dimensional surfaces applied to single particle analyses also provide evidence that the vRNA can be found within and outside of Gag condensates. ∩, colocalization (intersection); μm, micron.

### Addition of nuclear cell lysates induces large Gag-gRNA co-condensates

Based on our observation that the gRNA could dissolve Gag gels formed using crowding buffers (CB) (Figure 7), we examined whether whole cell lysates (WCL) could also promote formation of gRNA-Gag BMCs, as we have previously observed for HIV-1 NC protein [44]. Indeed, we observed that fluorescently labeled gRNA and Gag co-condensed and colocalized when CB was replaced with WCL derived from CD4^+^ T cells (Figure 8A), with the number and sizes of Gag-gRNA BMCs remaining similar to those produced by CB (Figure 8D, G).

**Figure 8:**
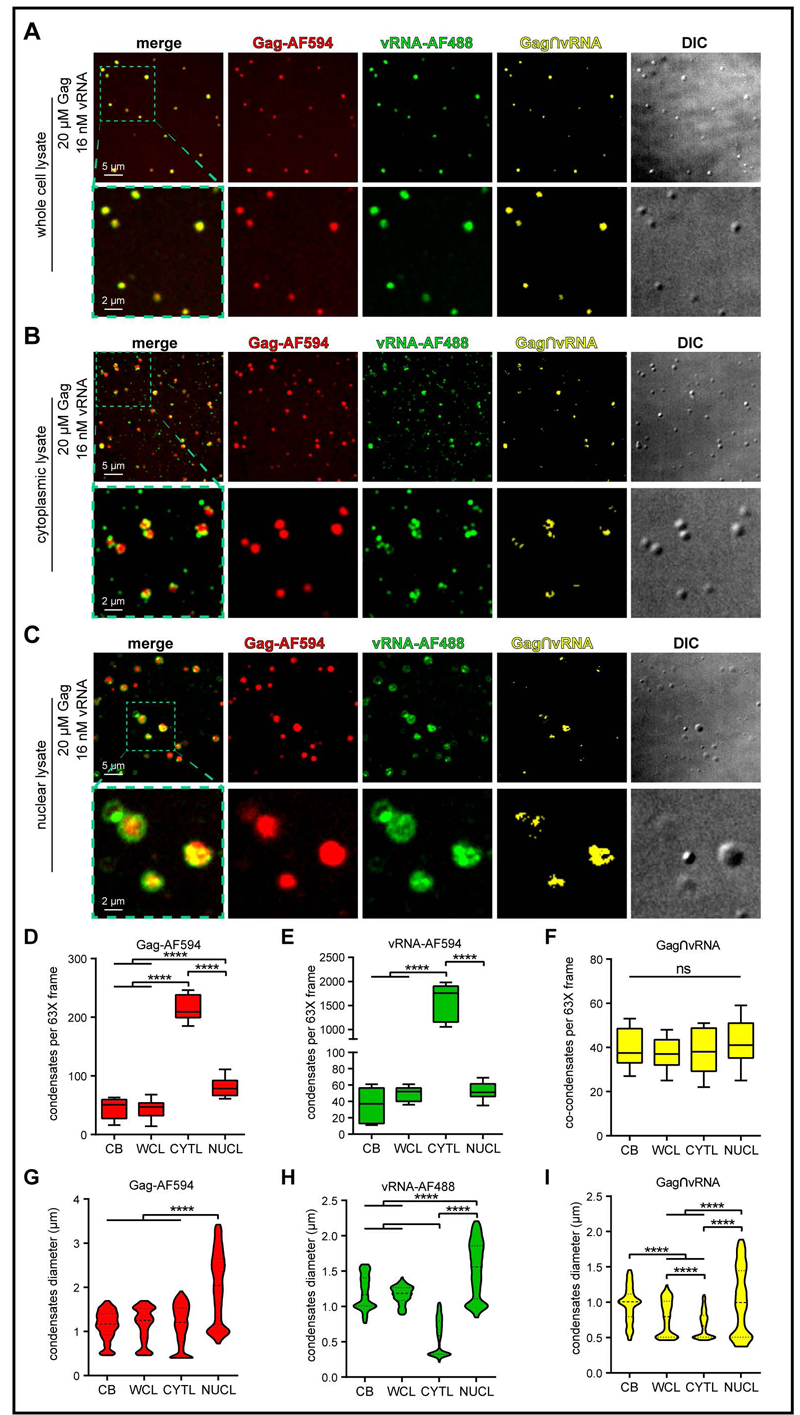
Nuclear and cytoplasmic lysates differentially influence Gag condensates. **(A)** Gag-AF594 and gRNA-AF488 phase separate into condensates when the crowding buffer is replaced with whole cell lysate from CD4^+^ T cells, as processed using NP-40 lysis buffer (150 mM NaCl, 10 mM Tris, 0.5% NP-40), where Gag and gRNA colocalizing condensates are visible. **(B)** Same as in (A), but where the crowding buffer is replaced with cytoplasmic lysate from nuclear-cytoplasmic fractionation. **(C)** Same as in (A), but where the crowding buffer is replaced nuclear lysate from nuclear-cytoplasmic fractionation. **(D-F)** Box and whisker plot corresponding to data from (A-C) and Figure 6. **(G-I)** Violin plots corresponding to sizes of condensates calculated from (A-C) and Figure 6. ****, p < 0.0001. CB, crowding buffer; WCL, whole cell extract; CYTL, cytoplasmic cell extract; NUCL, nuclear cell extract. ∩, colocalization (intersection); μm, micron.

Next, we examined whether there was a difference in Gag-gRNA BMCs formed by cellular constituents obtained from either cytoplasmic lysates (CYTL) or nuclear lysates (NUCL). To investigate this possibility, we performed nuclear and cytoplasmic fractionation of CD4^+^ T cells and used the same volume of buffers used for the WCL, taking into account the nuclear:cytoplasmic volumes of 5:1 for T cells to ensure accurate comparison to the WCL results [98]. Using the CYTL, we observed similar levels of co-condensation and colocalization, as well as similar sized BMCs, compared to CB or WCL (Figure 8B, G). However, CYTL increased the number of Gag condensates relative to the CB or WCL conditions (Figure 8B, D). We suspect that these gRNA condensates could be the result of cellular cytoplasmic RNA or RNA binding proteins forming complexes with the gRNA when mixed together. Unexpectedly, mixing Gag and gRNA with NUCL increased both the number of Gag-gRNA BMCs (Figure 8C, D), and strikingly, generated much larger condensates than observed in any other condition, with some as large as ∼4 microns, in contrast to ∼1 micron sizes observed using CB and WCL (Figure 8C, G). From past reports that nuclear RNAs induce solubility of condensates [96], it is possible that the higher abundance of differentially retained RNAs in the NUCL fraction [99, 100] may enhance the solubility of Gag-vRNA BMCs. The increased sizes of NUCL Gag condensates suggests that additional Gag and gRNA molecules were present in the complexes and/or that cellular factors were contained within individual BMCs, resulting in increased complex size. This result is consistent with our previous report showing that the nuclear Gag-viral RNA foci located at transcription sites were larger than viral ribonucleoprotein (vRNP) complexes at other cellular locations [91]. Additional studies are required to test the hypothesis that condensate-promoting nuclear host factors are present in the Gag-genomic RNA BMCs formed with NUCL. Another interesting observation was that CYTL also produced an increased number of smaller gRNA condensates devoid of Gag (i.e., 1364.12 ± 368.36 S.D.; p<0.0001), with a concomitant decrease in mean fluorescence intensity of colocalization between the gRNA and Gag (p<0.0001), (Figure 8B, C), despite no significance differences in the number of overall colocalizing gRNA and Gag condensates between NUCL and CYTL fractions (Figure 8F). In contrast, NUCL produced a higher number of Gag condensates devoid of gRNA (i.e., 27.05 ± 14.38 S.D. p<0.0001).

Finally, to examine whether the CTYL and NUCL differentially affected the position of the gRNA within condensates (Figure 8), we analyzed renderings of three-dimensional z-stacks for NUCL-and CTYL-induced co-condensates. In the case of the CYTL-induced condensates, we observed that the gRNA appeared in a circular pattern with focal regions and was located along the edges of Gag condensates (Figure 9A, B). When three-dimensional images of the smaller NUCL-generated co-condensates were analyzed, we observed that the gRNA was more condensed and located at the center of the condensates, resembling virion morphology (Figure 9C). In larger NUCL BMCs, the gRNA occupied a greater volume and colocalized with Gag along the perimeter of the droplets. In addition, there were smaller spherical protrusions of Gag-gRNA signal connected by a narrow stalk, suggesting that these could represent fission events from the larger droplets, supporting the idea that these condensates have liquid-like properties (Figure 8C) [25].

**Figure 9:**
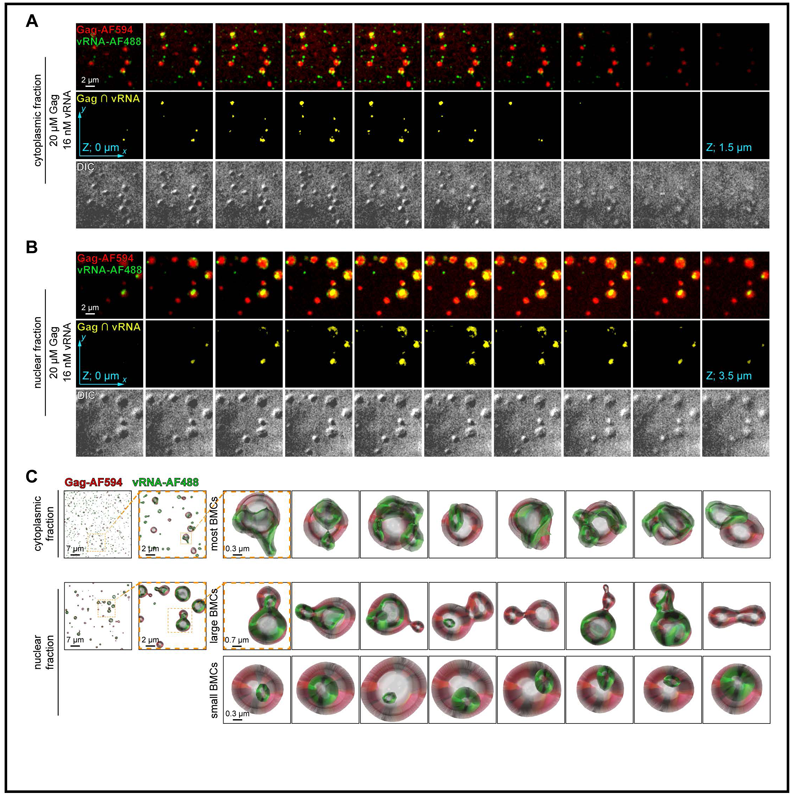
Nuclear lysates provide increased liquidity to Gag condensates. **(A)** Analyses of Z-stacks Gag-AF594 and gRNA-AF488 positive condensates when the crowding buffer is replaced with Jurkat T cell cytoplasmic lysates from nuclear-cytoplasmic fractionation, as processed using NP-40 lysis buffer (150 mM NaCl, 10 mM Tris, 0.5% NP-40). Gag and gRNA colocalizing condensates are visible and colocalization is mostly central to Gag condensates. **(B)** Same as in (A), but where the crowding buffer is replaced with Jurkat T cell nuclear lysates from nuclear-cytoplasmic fractionation. **(C-D)** Three-dimensional surfaces applied to single particle analyses show that the gRNA can be found within and outside of Gag biomolecular condensates (BMC) when cytoplasmic lysates are applied (C), and is mostly found inside of both smaller and larger condensates produced by nuclear lysate generated BMCs. ∩, colocalization (intersection); μm, micron.

## Discussion

In this study, we performed *in vitro* studies using purified HIV-1 Gag proteins to determine whether IDR-containing MA and NC domains of Gag promote its phase separation. We found that the MA and NC domains of Gag differentially effected the formation of Gag condensates. At the higher protein concentrations tested, WT Gag produced a gel. None of the mutations tested abolished the ability of the protein to form BMCS but only altered the conditions under which BMCs formed. Mutation of the MA basic residues produced smaller condensates, and NC ZF mutants formed larger condensates, suggesting that both the NC and MA domains of Gag participate in the formation of Gag-gRNA BMCs (Figures 2-4). Although we examined MA mutants within the context of full-length Gag, in mature viruses MA is cleaved from precursor pr55Gag during virus maturation, post virion release, to form a discontinuous matrix shell that lacks order and is associated with the inner leaflet of the viral membrane [101], not with the gRNA. Our results are the first to provide evidence that MA also promotes Gag condensation when NC ZnFs are mutated. Just as Gag is found to self-assemble *in vitro* as rod-like structures at lower pH [102], we observed a similar shape for some Gag BMCs at different experimental conditions tested (Figure 2-4). Larger spherical and rod-shaped BMCs formed by Gag MA mutants at higher protein and lower salt concentrations suggests a mechanism by which MA may assist the conformational switch from MA-NC contacts in a hairpin conformations to an extended conformation found in virions [102]. To provide insights from this biophysical study, we have compiled a table summarizing how the different Gag MA and NC mutants tested herein affect the budding and gRNA binding activities of Gag in cells (Table 1).

HIV-1 gRNA induces Gag dimerization [103], whereas Gag multimerization requires the NC domain and is promoted by CA dimers and the MA RNA binding domain (MA-RBD) [104]. In this report, we provide evidence that at low protein concentrations, gRNA induced the formation of larger Gag condensates (Figure 5-6). Interestingly, these Gag-gRNA complexes were both larger and more spherical compared to those formed by NC Zn mutants mixed with gRNA, unlike the MA mutants (Figure 6). At lower protein concentrations, MA mutants produced fewer and smaller spherical aggregates, suggesting that MA was important for maintenance of sphericity of BMCs [105]. Less spherical condensates were also observed at increased salt and protein concentrations for the MA deletion mutant, accompanied by a shift in gel formation at the highest protein and salt concentrations (Figure 4). Conversely, at higher protein concentrations, we observed that Gag NC ZF mutants generated much larger condensates with decreased sphericity (e.g., especially evident for ZnF1 and ZnF1/2 NC mutants) (Figure 2, 3), suggesting that this domain confers a less liquid-like state to Gag condensates. We previously reported that 1,6-hexanediol readily dissolves HIV-1 NC protein condensates that otherwise can grow into large SGs [44, 60]. Since neither HIV-1 Gag nor CA forms large aggregates in cells, it is possible that the restrictive MA domain is responsible for chaperoning the gRNA-bound NC condensates until a sufficient number of Gag molecules are present for initiation of higher order structures [103].

Nuclear residency of Gag has been reported in a number of studies [106-109], interacting with nuclear host factors [106, 110] and binding to unspliced HIV-1 RNA at discrete sites within the perichromatin space [91]. In this work, we observed that the gRNA-Gag condensates formed with nuclear extracts were larger than their cytoplasmic counterparts (Figure 7), consistent with being more liquid-like and could contain additional cellular proteins or nucleic acids. This phenomena may be mirrored by liquid-like condensates formed with RNA-binding proteins with nuclear function and origin (e.g., FUS, TDP43, and hnRNPA1) which, when mutated and/or impaired in their nucleocytoplasmic shuttling [111], are re-localized to the cytoplasm where they form solid aggregates [112, 113] leading to neurodegenerative diseases [10, 11]. The formation of larger, possibly more liquid-like Gag BMCs induced by the RNA-rich nuclear environment may ensure that the Gag-gRNA complex remains intact as it traverses the nuclear envelope, encounters the cytoplasmic environment which could impede its transport, and ultimately reaches the plasma membrane. When mixed with cytoplasmic lysates, we observed an increased number of cytoplasmic Gag BMCs lacking gRNA. These Gag BMCs could be associated with other cytoplasmic RNAs or proteins and involved in different functions other than gRNA packaging. Taken together, these observations suggests that a nuclear origin may promote the specific packaging of gRNA into Gag BMCs. Although oligomerization of Gag mediated by CA and NC would generate a large viral ribonucleoprotein complex that would need to cross the nuclear envelope, recent observations suggest that the large, incoming intact pre-integration complex is capable of transit through the nuclear pore [114, 115].

Altogether, these results are consistent with past observations of mechanistic properties of both viral and non-viral phase separating BMCs and warrant future investigation into how HIV-1 BMCs participate in the biogenesis of the growing virion during the assembly process. Such studies could uncover new ways to target viruses like HIV-1 that utilize disordered protein elements for their condensation into vRNPs, enabling them to traverse the intracellular environment from nucleus, through the cytoplasm, to the plasma membrane. The current views of nuclear pore flexibility [116, 117], along with evidence that many viral components contain disordered domains and form BMCs [18, 29] has the potential to replace virion rigidity dogmas with a new view of viruses as flexible shapeshifters which evade the immune system and are less vulnerable to antibody detection and neutralization as a result of their disorder [38].

**Table 1 in separate document**

## RESOURCE AVAILABILITY

### Materials Availability

Further information and requests for resources, code and reagents should be directed to and will be fulfilled by. There are restrictions to the availability of reagents reported in this study due to the lack of an external centralized repository for its distribution and our need to maintain the stock. We are glad to share reagents with reasonable compensation by requestor for its processing and shipping.

**Table 1.**

| **Gag** | **Budding** | **RNA Binding** | **Correlative Condensate Features (this study)** |
| --- | --- | --- | --- |
| MA 7N | Redistributes Gag localization, increasing cytoplasmic localization at perinuclear regions [118]; decreased membrane binding capacity and overall Gag production, suggesting Gag instability [52]. | Substitution of 2 or more basic residues abolishes RNA binding [119]; 7N mutant decreased binding affinity to monomeric and dimeric psi RNA and gRNA packaging enhancer (GRPE) region [120, 121]. | Condensates less spherical at optimum protein concentrations; rod-like at high protein concentration; Gag-gRNA BMCs less spherical at low protein concentration and little impact from gRNA on BMC morphology and decreased gelling at higher protein concentrations. |
| MA Δ15-39 | Decrease in budding and stability of Gag [52]. MA N-terminal region residues K26 and K27 required for nuclear accumulation [122]. | Deletions of MA basic residues eliminates RNA binding [119]. | Increased self-association of Gag; more severe loss of sphericity at optimal protein concentrations; rod-like at high protein concentration; Gag-gRNA BMCs less spherical at low protein concentration and decreased gelling at higher protein concentrations. |
| NC ΔZF1 | Delayed Gag-gRNA trafficking to the plasma membrane [90]. Increased assembly time of Gag at the plasma membrane [123], but does not hinder VLP production [124]. | Decreases specific recognition of psi RNA [87]. Does not affect Gag cytoplasmic binding to gRNA [90]. | Decreased condensate formation at optimum protein concentrations; increased particle size and decreased sphericity at high protein concentration; low Gag-gRNA BMCs heterogeneity at low protein concentration and notable decrease in condensates and further loss in sphericity at higher protein concentrations. |
| NC ΔZF2 | More prominently delayed Gag-gRNA trafficking to the plasma membrane compared to ΔZNF1 [90]. Increased assembly time of Gag at the plasma membrane [123]. | Does not affect Gag cytoplasmic binding to gRNA [90]. | Decreased condensate formation and rod-like at near optimum protein concentrations; increased particle size at high protein concentration; low Gag-gRNA BMCs heterogeneity at low protein concentration and notable decrease in condensates sphericity at higher protein concentrations. |
| NC ΔZF1/2 | Increased assembly time of Gag at the plasma membrane [123] and decreased VLP production [124]. | Results in severe decrease of Gag binding to psi [125]. | Decreased condensate formation at optimum protein concentrations; largely increased particle size and decreased sphericity at high protein concentration; low Gag-gRNA BMCs heterogeneity at low protein concentration and notable decrease in condensates sphericity at higher protein concentrations. |

### Data and Code Availability

- Data reported in this paper will be shared by contacts upon request.
- This paper does not report original code.
- Any additional information required to reanalyze the data reported in this paper is available from the contacts upon request.

### Experimental Model and Subject Details

#### Cells and cell culture

Adherent HEK293T cells originated from a human female embryonic kidney. HEK293T cells [American Type Culture Collection (ATCC)] were grown and maintained in Dulbecco’s Modified Eagle Medium (DMEM) (GIBCO Thermo Fisher Scientific #) supplemented with 10% fetal bovine serum (FBS) (Hyclone), 1% penicillin/streptomycin

(Invitrogen) at 37°C and 5% CO_2_. The CD4^+^/CXCR4^+^ Jurkat CE6.1 T cell line (ATCC) originates from peripheral blood and was grown and maintained as suspension culture in RPMI 1640 (Life Technologies) supplemented with 10% FBS (Hyclone) and 1% penicillin/streptomycin (Life Technologies) at 37 °C and 5% CO_2_.

### Method Details

#### Cloning and plasmids

The various pET28a(-His) HIV Gag mutants were cloned using the Q5 site-directed mutagenesis kit according to the manufacturer’s instructions (NEB) and primers were designed using NEBaseChanger program (nebasechanger.neb.com). A full-length pET28a(-His) HIV Gag WT construct [126] was modified to contain the mutations described below. The pET28a(-His) HIV Gag MA 7N mutant was generated using primers 5’- AGG GGG AAA CAA CCA ATA TAA CCT AAA ACA TAT AGT ATG GGC AAG C-3’ and 5’- GGA TTT AAG TTA ATA TTT TCC CAG TTA TCT AAT TCT CCC CCG CT-3’. The pET28a(-His) HIV Gag MA Δ15-39 mutant was generated using primers 5’- GAG CTA GAA CGA TTC GCA G-3’ and 5’- ATC TAA TTC TCC CCC GCT-3’. pET28a(-His) Gag NC.ZnF1 mutant was cloned using primers 5’- GGG CAC ATA GCC AAA AAT AGC AGG GCC CCT AGG AAA AAG-3’ and 5’- TTC TTT GCC ACT ATT GAA ACT CTT AAC AGT CTT TCT TTG GTT CC3’. pET28a(-His) Gag NC.ZnF2 was generated using primers 5’- GGA CAC CAA ATG AAA GAT AGT ACT GAG AGA CAG GCT AAT TTT TTA G-3’ and 5’- TTC CTT TCC ACT TTT CCA ACT GCC CTT TTT CCT AGG GGC-3’. The pET28a(-His) Gag NC.ZnF1 & ZnF2 double mutant was generated by mutating the second CH box of pET28a(-His) Gag NC.ZnF1 using the same primers used to clone pET28a(-His) Gag NC.ZnF2.

#### Recombinant HIV-1 Gag proteins

A full-length recombinant HIV-1 NL4-3 pr55 Gag protein, stored in 50 mM Tris-HCl, 1 M NaCl, pH 8.0., was acquired from the NIH AIDS Reference and Reagent Program (ARRP) (ARP-13385; cat#13385, lot#190002). For expression and purification of other WT and mutant recombinant HIV-1 Gag proteins used in this study, BL21(DE3) pRIL cells containing the appropriate HIV Gag T7 expression plasmid were inoculated from an 8% glycerol MDG stock by scratching cells from the top layer with a sterile pipet and dipping the pipet into a test tube containing 2mL of MDG or MDAG-135 non-induction media [127] supplemented with 15 μg/mL chloramphenicol and 200 μg/mL kanamycin. These cultures were grown overnight at 37°C in a floor shaker (250 rpm). The following day the overnight culture was used to inoculate (1:500) 250mL of ZYM-or ZYP-5052 autoinduction media supplemented with 15μg/mL chloramphenicol and 100μg/mL kanamycin. The autoinduction culture was grown for 9-12 hrs at 37⁰C until it reached saturation. Longer growth times lead to cell lysis and substantial reduction of the yield and purity of Gag. The cells were harvested by centrifugation for 15 min at 6000xg and 4°C. The media was removed, and the cell pellets were drained by inversion over a tissue for 5min. The cell pellets were either used immediately or frozen and stored at −20°C.

Wild-type HIV Gag-His-Tev and the non-tagged HIV Gag variants were purified using a modification of our previously described protocol [126]. Briefly, cells (3-5 g) were resuspended in 50 mL of ice-cold cell lysis buffer (50mM Tris·HCl pH 8.0; 500 mM NaCl; 20mM Imidazole; 0.1mM TCEP; 0.1mM EDTA; 0.1% NP-40 (Tergitol), 10% glycerol, and 2 tablet Roche Complete Protease inhibitor cocktail). The buffer was supplemented with 30 kU/gm cell paste rLysozyme (Millipore) and then homogenized in a Dounce homogenizer to remove any cell clumps. The cell suspension was lysed by three passages through an MP-120 microfluidizer using 12000 psi during the first two passages and 27000 psi for the final passage. The cell debris was removed by centrifugation at 40,000 *g* for 30 min. The supernatant was transferred to a small beaker, and the nucleic acids were precipitated at 4°C by slowly adding PEI from 10% (w/v) pH 7.8 stock to a final concentration of 0.14%. This solution was allowed to stir on ice for 15min, and the precipitate was removed by centrifugation at 40,000 *g* for 10 min. The supernatant was transferred to a beaker, and the protein precipitate by the addition of a room temperature saturated ammonium sulfate stock pH 7.0 to a final concentration of 30% (v/v) saturation. The solution was stirred on ice for 30 min, and the precipitate containing Gag was collected by centrifugation at 40,000 *g* for 30 min. The residual ammonium sulfate solution was removed by inverting and draining the tubes over a kimwipe (Kimberly-Clark) for 10 min. This pellet contains the majority of the Gag in a highly enriched form.

For HIV Gag-His-Tev the pellet was resuspended directly in 50mL of ice-cold Ni-A buffer (50 mM HEPES pH 7.5; 500 mM NaCl; 20 mM Imidazole; 0.1 mM EDTA, 0.1 mM TCEP, and 2 tablet Roche Complete Protease Inhibitor Cocktail). Any residual insoluble material was removed by centrifugation at 40,000 *g* for 15 min, and the supernatant was then loaded onto a 5 mL His-Trap FF crude column equilibrated in Ni-A. The column was washed with 95% Ni-A and 5% Ni-B (50 mM HEPES pH 7.5; 500 mM NaCl; 400 mM Imidazole; 0.1 mM EDTA, 0.1 mM TCEP) and the bound HIV-Gag-His-TEV eluted in a 100% Ni-B step gradient. The peak fractions were collected, and the concentration of the eluted protein was estimated using a nanodrop spectrophotometer from the absorbance at 280 nm. To remove the His tag from Gag, TEV protease was added to the pooled peak fraction at a concentration of (25:1 Gag:TEV), and the sample was then dialyzed against 2L of Dialysis buffer (25 mM HEPES pH 7.5; 500 mM NaCl; 60 mM Imidazole; 0.1 mM EDTA and 0.1 mM TCEP) overnight at 4⁰C. After dialysis, the protein was harvested and loaded onto a 1 mL His-Trap ff crude Ni-Affinity column equilibrated in Dialysis buffer. The His-tag removed Gag elutes in the flow through from this column, and TEV, uncleaved Gag, and the His_6_ peptide are eluted in a 100% Ni-B bump of this column. The flow through from the second Ni-affinity column was diluted with ice-cold Dilution buffer (10 mM HEPES pH 7.5; 0.1 mM EDTA and 0.1 mM TCEP) to a final salt concentration of ∼200 mM NaCl and loaded onto a 5 mL Hitrap SP ImpRES ion exchange column equilibrated in SP-A buffer. The column was eluted in a 20 column volume linear gradient from 100% SP-A to 25% SP-A:75% SP-B buffer (10 mM HEPES pH 7.5; 2 M NaCl; 0.1 mM EDTA and 0.1 mM TCEP). At this point, the protein is highly pure and dialyzed against 3 changes of Storage-500 buffer (10 mM HEPES pH 7.5; 500 mM NaCl; 0.1 mM EDTA; 0.1 mM TCEP; 0.01 mM ZnCl_2_). The purified HIV Gag-Tev protein was then concentrated to 2-4mg/mL using a 30k Centricon (Amicon) centrifugal filter, aliquoted and snap-frozen in a dry ice ethanol bath, and stored at −80⁰C until it was used. In some instances, the peak fractions from the SP column were concentrated to 2-4 mg/mL and further purified by chromatography on a Superdex S200 10x300 (Cytiva) column equilibrated in Storage-500 buffer. The peak fractions from this column were pooled and then concentrated to 2-4 mg/mL using a 30k cutoff Amicon centrifugal filter. The concentrated protein was frozen and stored at −80⁰C, as before, until used. The addition of the size exclusion purification step did not markedly increase the purity of the protein and led to sizable losses due to the significant tailing of the eluted peak and increased losses due to the additional concentration step.

For the untagged HIV Gag variants, the cell pellets were resuspended in 20mL of ice-cold buffer comprised of 10 mM HEPES pH 7.5; 500 mM NaCl; 0.1 mM EDTA, and 0.1 mM TCEP, and then diluted to 50 mL with ice-cold dilution buffer containing 1 Roche Complete protease inhibitor cocktail tablet. The sample was centrifuged at 40kxg for 15 minutes and then loaded onto a 5 mL HiTrap SP ImpRes column equilibrated and in SP-A. The column was eluted using the same program used for HIV Gag-Tev above. The peak fractions were diluted 1:1 with SP-A buffer and loaded onto a 5 mL HiTrap MMC (Cytiva) column equilibrated in this buffer. The bound protein was then eluted in a 20CV linear gradient from 100% SP-A buffer to 75% MMC buffer (10 mM HEPES pH 7.5; 500 mM Arginine; 1 M NaCl; 0.1 mM EDTA and 0.1 mM TCEP). The peak fractions were then pooled and concentrated to 5-10mg/mL and further purified by chromatography on a Superdex 200 10x300 (Cytiva) column equilibrated in Storage-400 buffer (10 mM HEPES pH 7.5; 400 mM NaCl; 0.1 mM EDTA; 0.1 mM TCEP and 0.01 mM ZnCl_2_. The peak fractions were concentrated to 4-6 mg/mL by ultrafiltration using a 30k cutoff Amicon ultrafiltration device, snap-frozen, and stored at −80⁰C until used. All recombinant proteins were resuspended in 10 mM Hepes pH 7.5, 500 mM NaCl, 0.1 mM TCEP, 0.1 mM EDTA, 20 uM ZnCl_2_, with exception of the NC ZnF2 mutant in 10 mM Hepes pH 7.5, 1.0 M NaCl, 0.1 mM TCEP, 0.1 mM EDTA, 0.01 mM CaCl_2_.

#### Viral RNA extraction and labeling

For virus production and isolation of WT vRNA, HEK293T cells (1 × 10^6^) were plated in 10-cm dishes in DMEM supplemented with 10% FBS at 37°C and 5% CO_2_ for 24 h, and were then transfected with 10 μg of pNL4-3 DNA for 6 h. The supernatant was collected 24 h later, spun down at 3,000 rpm for 20 min, filtered by a 0.45 µm filters, and then spun at 20,000 rpm for 1.5 h. 50 µL D-PBS was used to resuspend the virus pellets, which were stored at −80°C until further use. For viral RNA extraction, 200 µL of virus solution in D-PBS was mixed with 1 mL Trizol and 200 µL of chloroform before being spun down at 12,000 rpm for 15 min. The aqueous phase was transferred to a new tube and incubated with 500 µL isopropanol for 10 min at room temperature, then spun at 12,000 rpm for 10 min at 4°C. The supernatant was discarded, and the vRNA pellet was washed using 1 mL of RNase-free 75% alcohol, and spun down at 7500 *g* for 5 min at 4°C. The supernatant was discarded, and the purified vRNA was resuspended in 50 µL RNase free H2_O_. vRNA concentration was determined using a NanoDrop ND-100 spectrophotometer (ThermoFisher Scientific).

For fluorescent vRNA experiments, vRNA was labeled using fluorescence *in situ* hybridization as previously described [128, 129], where vRNA was fixed with 4% paraformaldehyde for 20 min in D-PBS (Wisent) containing RNAseOUT (Invitrogen), and was then incubated with a digoxigenin-labeled RNA probe synthesized in vitro in presence of digoxigenin-labeled UTP (Roche) for 30 mins at 42°C, followed by incubation with sheep anti-digoxigenin-AP Fab fragments (Sigma) for 30 mins at 37 °C, and a final incubation with Donkey anti-Sheep IgG (H+L) Cross-Adsorbed Secondary Antibody, Alexa Fluor® 488 (Invitrogen) for 30 mins at 37 °C.

#### Creation of phase separated condensates

Formation of *in vitro* phase separated condensates was monitored by differential interference contrast (DIC) and fluorescence microscopy as previously described [13, 44, 45]. For examination of fluorescent condensates, purified proteins were labeled using the Alexa Fluor™ 594 Microscale Protein Labeling Kit (Thermo Fisher Scientific, #A30008) according to manufacturer’s instructions. Recombinant proteins were mixed with buffers containing 20 mM HEPES (VWR; cat#CA-EM5320), varying concentrations of NaCl, pH 7.4, and with 150 mg/ml ficoll (Lymphocyte Separation Medium, Corning; cat#25-072-CV) added as molecular crowding agent. For imaging of *in vitro* condensates, 6 μL of sample mixtures were loaded onto 25 x 75 mm x 1mm thick glass slides (Thermo Scientific), and covered with 18 mm ø No. 1 cover glasses (VWR VistaVision^TM^, VWR International), and were sealed with clear enamel (Revlon).

#### Cell fractionation

Cell fractionation was performed by collecting and washing 5 x 10^7^ cells three times with 4°C PBS at 1500 rpm for 10 mins, which was discarded and replaced with cytoplasmic lysis buffer (10 mM KCl, 0.1 mM EDTA, 10 mM HEPES, pH 7.9. 0.4% NP-40 (IGEPAL-30) with and DNAseI for 10 mins on ice, followed by centrifugation at 18,100 rcf for 10 minutes at 4°C. Supernatants were transferred to new sterile 1.5 mL microtubes, and the remaining nuclear pellet was washed three times with 4°C PBS at 1500 rpm for 10 mins, which was discarded and replaced with cytoplasmic lysis buffer [10 mM KCl, 0.1 mM EDTA, 10 mM HEPES, pH 7.9. 0.4% NP-40 (IGEPAL-30)] with DNAseI for 10 mins on ice, ahead of homogenization using QIAshredder columns (QIAGEN), used as recommended by manufacturer. The resulting lysate was centrifuged at 18,100 *g* for 10 minutes at 4°C.

#### Imaging and analyses of phase separated condensates

Condensates were observed by microscopy 10 mins later, for both fluorescence coinciding with clearly discernible spheres using DIC. This was performed using a Leica DM16000B microscope equipped with a WaveFX spinning disk confocal head (Quorum Technologies) and 63X, Oil objective, where images were acquired with a Hamamatsu EM-charge coupled device digital camera. Single-slice and Z-stack scanning was performed and digitized at a resolution 1,024 × 1,024 pixel. For multi-color image capture, blue and yellow-green excitation lasers were used to excite AlexaFluor-594 and 488, and emissions were sequentially captured with 570–620 and 500–550 nm bandpass filters, followed by differential interference contrast (DIC) image capture. Raw .liff files were exported by the Volocity software (Perkin Elmer) for import into the Imaris and ImarisColoc software v.9.7.2 (Andor) used for generation of new colocalization channels, and .csv exports of quantitative measurements of mean signal intensity values used for downstream data harmonizing and statistical analyses using Excel (Microsoft) and GraphPad Prism v.8.0.1, respectively.

#### Microscopy and imaging analyses

Laser confocal microscopy was performed using a Leica DM16000B microscope equipped with a WaveFX spinning disk confocal head (Quorum Technologies) and 63X, Oil objective, and images were acquired with a Hamamatsu EM-charge coupled device digital camera. Scanning was performed and digitized at a resolution 1,024 × 1,024 pixel. For multi-color image capture, AlexaFluor-594 and -488, emissions were sequentially captured with 570–620 nm, and 500–550 nm, bandpass filters, followed by DIC image capture.

#### Informatics

DisEMBL 1.5 predictor of intrinsic protein disorder (http://dis.embl.de/), as well as a Predictor of Natural Disordered Regions (PONDR; http://www.pondr.com/) were used to analyze protein disorder from protein FASTA sequences obtained from the NCBI protein sequence database.

#### Quantification and Statistical Analyses

All experiments were performed in triplicate with similar results, unless otherwise indicated in figure legends. Three independent observers validated phenotypes resulting from all experimental conditions tested. Condensate imaging statistics result from counting an average of *n* = 2,000 per condition tested. Data analysis and plotted graphs were performed using GraphPad Prism v.8.0.1, and one-way ANOVA (with Tukey’s multiple-comparisons test) and 95% CI was used for multiple comparisons. P-values <0.05 were considered to indicate a statistically significant difference. Box-plot center lines indicate median, box limits are interquartile range and whisker are minimum to maximum. Violin plots show min–max ranges, with center dashed lines indicating median, dotted lines as interquartile ranges, and kernel densities as distributions of each group.

## Acknowledgements

We thank lab members from all collaborating laboratories for comments on the work presented in this manuscript; Mathew Duguay and Christian Young of the LDI cell imaging and flow cytometry core facility for assistance with microscopy; and the NIH Reference and Reagent Program for antibodies and reagents. We thank Malgorzata Sudol, MS, for technical assistance. Research reported in this publication was supported by the National Institute On Drug Abuse of the National Institutes of Health under Award Number R21DA053689 to LJP with subaward to AJM. The content is solely the responsibility of the authors and does not necessarily represent the official views of the National Institutes of Health. This work was also supported by grants from the Canadian Institutes of Health Research (CIHR) (FRN-162447 to A.J.M. and FRN-PJT-178165 to A.W.C.).

## Author Contributions

AM, LJP and AJM conceived of the study, designed experiments, interpreted data; AM and MN performed the experiments; JMF and RKM performed cloning, protein purification, data interpretation, experimental design, and writing; GSL, AC, and JC - experimental design, data interpretation, and writing; LJP and AJM supervised and obtained funding; and AM drafted the manuscript and all authors revised and approved the final version.

